# Low-complexity domains adhere by reversible amyloid-like interactions between kinked β-sheets

**DOI:** 10.1101/153817

**Authors:** Michael P. Hughes, Michael R. Sawaya, Lukasz Goldschmidt, Jose A. Rodriguez, Duilio Cascio, Tamir Gonen, David S. Eisenberg

## Abstract

Control of metabolism by compartmentation is a widespread feature of higher cells. Recent studies have focused on dynamic intracellular bodies such as stress granules, P-bodies, nucleoli, and metabolic puncta. These bodies appear as separate phases, some containing reversible, amyloid-like fibrils formed by interactions of low-complexity protein domains. Here we report five atomic structures of segments of low-complexity domains from granule-forming proteins, one determined to 1.1 Å resolution by micro-electron diffraction. Four of these interacting protein segments show common characteristics, all in contrast to pathogenic amyloid: kinked peptide backbones, small surface areas of interaction, and predominate attractions between aromatic side-chains. By computationally threading the human proteome on three of our kinked structures, we identified hundreds of low-complexity segments potentially capable of forming such reversible interactions. These segments are found in proteins as diverse as RNA binders, nuclear pore proteins, keratins, and cornified envelope proteins, consistent with the capacity of cells to form a wide variety of dynamic intracellular bodies.

**One Sentence Summary:** Atomic structures show transient membraneless organelles of cells formed by a new type of protein interaction akin to pathogenic amyloid fibrils.

## Main Text

The human proteome is replete with domains having highly overrepresented subsets of amino acids. These are termed Low-Complexity (LC) domains (Fig. 1A and B). LC domains are often intrinsically disordered (*1*), and they are dramatically underrepresented in the Protein Data Base (PDB) of known 3D structures (*2*). In the tree of life, higher organisms typically have more proteins with LC domains: 4% of proteins in *Escherichia coli*, 18% of proteins in *Saccharomyces cerevisae*, and 29% of proteins in *Homo sapiens* have a LC domain. The role of the LC domains found in higher organisms has recently been illuminated through studies of RNA-binding proteins: these domains aid in the transient formation of functionally important intracellular compartments (*3*–*8*).

**Fig. 1.**
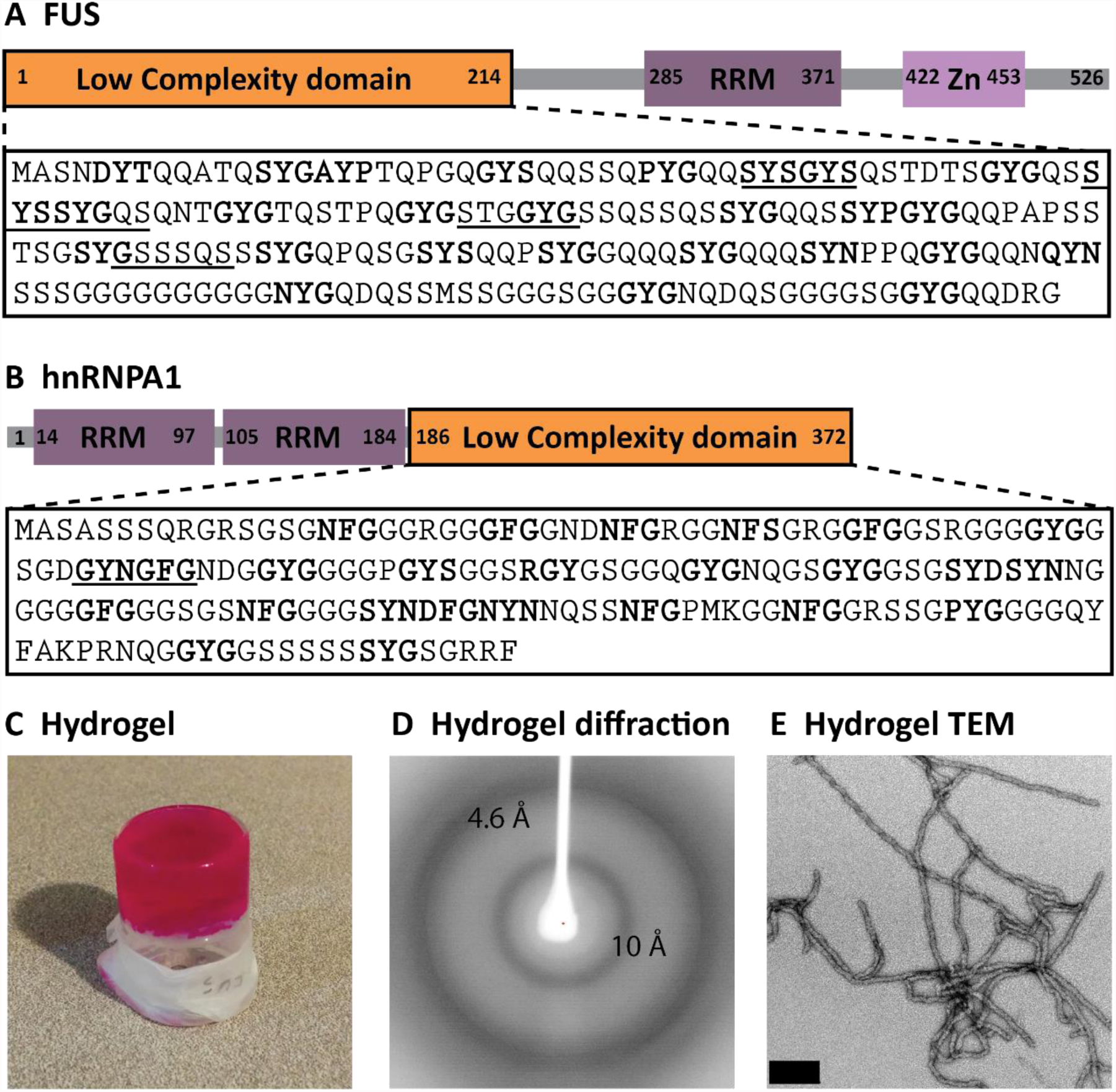
Low complexity (LC) domains of FUS and hnRNPA1. (**A** and **B)** Amino acid sequences of the LC domains of FUS and hnRNPA1. Residues [G/S]Y[G/S] motifs are in bold and sequences of structures determined in this work are underlined. **(C)** Hydrogel formed from the LC domain of FUS fused with mCherry. The hydrogel sits atop a 5 ml beaker which had previously been inverted. **(D)** X-ray fiber diffraction pattern of the mCherry-FUS hydrogel, displaying the cross-β diffraction pattern characteristic of amyloid structure. **(E)** Negatively stained transmission electron micrograph of 400x diluted mCherry-FUS hydrogel shows amyloid-like fibrils. Scale bar is 0.2μm.

In mammalian cells, RNA binding proteins are enriched in LC domains (*3*). RNA binding proteins are commonly found in dynamic intracellular bodies, composed of RNA and protein. These bodies are non-membrane bound organelles that appear and disappear in response to stimuli; they include P-bodies, nuclear paraspeckles, and stress granules (SGs). The LC domains of RNA binding proteins have been shown to localize them to SGs and nuclear paraspeckles (*3*, *4*), suggesting that the function of many LC domains is to organize dynamic intracellular bodies in the cell.

From a structural and biophysical perspective, one wants to know the nature of the adhesive interactions between LC domains that guide the formation of these dynamic intracellular bodies. From in vitro studies of purified LC domains, two states have been observed: a liquid-liquid phase separation and a solid phase hydrogel. At concentrations comparable to those found in cells, the SG associated proteins hnRNPA1, hnRNPA2, and FUS undergo liquid-liquid phase separation (*9*–*12*). Liquid-liquid phase separation is a general property of macromolecules that have multivalent interactions with each other, causing them to separate into two liquid phases, one having ^~^100 times the concentration of the macromolecule compared to the bulk phase (*13*). Over time, or at higher concentrations of protein, the LC domains of hnRNPA1, hnRNPA2, FUS, and RBM14 can transition into a reversible solid-phase hydrogel (*3*, *4*, *6*, *9*, *12*). Electron microscopy reveals that such hydrogels are composed of protein fibrils, and X-ray diffraction of the hydrogel yields an amyloid-like cross-β pattern (Fig. 1C-E) (*3*, *14*). This implicates amyloid formation as a possible basis for the structures formed from LC domains.

Amyloid fibrils contain pairs of closely mating β-sheets along the fibril axis. Residue side-chains extend across this interface to tightly interdigitate with sidechains of the opposing β-sheet to form an extensive, dry interface, as seen in the structure of NKGAII from amyloid-beta (Aβ) (Fig. 2F). This structure is called a steric zipper, and it forms the spine of amyloid fibrils (*15*, *16*). The steric zipper explains the extraordinary stability of some pathogenic amyloid fibrils that are resistant to denaturation by SDS and boiling. This contrasts starkly with the amyloid-like fibrils found in hydrogels formed from the LC domains of FUS, which are SDS-sensitive (*3*). The lability of FUS amyloid-like fibrils may be central to the dynamic nature of SGs.

**Fig. 2.**
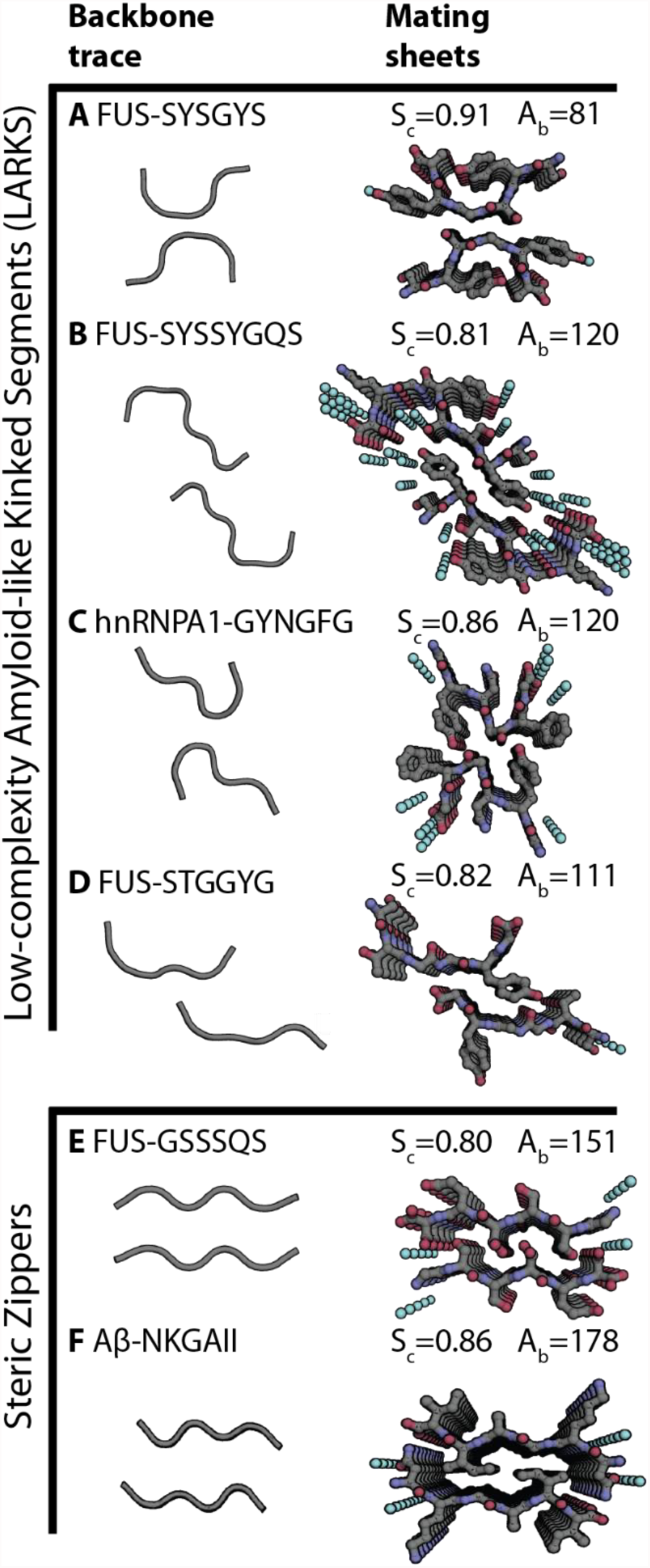
Structures of LARKS (A-D) compared to steric zippers (E-F). All structures are viewed down the fibril axis of two mated β-sheets. The left-hand column shows the trace of the backbones of mated sheets to highlight kinks in the backbones of LARKS and the pleating of the classical β-sheets in steric zippers. The right-hand column shows the atomic structures of mated sheets. Each interface is characterized by the steric complementarity (S_c_) score and buried surface area (A_b_) in Å^2^ between the mated sheets. Carbon atoms are colored gray, Nitrogen is blue, Oxygen is red, and water molecules are shown as aqua spheres. Five layers of β-sheets are shown of the hundreds of thousands in the crystals.

The adhesive interactions between LC domains must be both stabilizing enough to ensure their formation and weak enough to be reversible. Fluorescence measurements of SGs in cells indicate a rapid turnover of molecules in a SG, similar to that found in liquid-liquid phase separations (*11*, *17*). However at the core of these granules is a more solid phase state (*18*). Mass spectrometry footprinting of hnRNPA2, either extracted from cells or from hydrogels formed in vitro, shows similar patterns (*12*). Taken together, these experiments suggest that the amyloid structures formed by LC domains are labile, and therefore may form the basis for dynamic intracellular bodies. Here we present atomic structures of segments from the LC domains of hnRNPA1 and FUS that offer insight into how LC domains can form fibrils, yet remain reversible, in contrast with pathogenic amyloid fibrils.

To understand the adhesive interactions between LC domains of proteins recruited to SGs, we sought relevant atomic structures. We were guided by the studies of the LC domains FUS and RBM14, which show that successive replacement of tyrosine residues by serine lower their capacity to form hydrogels (*3*, *4*). Accordingly we scanned the LC domain of FUS for tandem sequence motifs of the form [G/S]Y[G/S], finding two such segments: FUS-^37^SYSGYS^42^ (SYSGYS) and FUS-^54^SYSSYGQS^61^ (SYSSYGQS) (Fig. 1A). Confirming the relevance of these crystals to adhesive interactions of LC domains, both segments crystalized as micron-sized needles, from which powder X-ray diffraction reproduced the 4.65 Å ring and other features observed by diffraction of the FUS-LC hydrogel (fig. S1). Atomic structures of both segments were determined, in addition to the structures for three other segments, which were identified by 3D profiling (see below). These are: ^243^GYNGFG^248^ (GYNGFG) from the protein hnRNPA1 (Fig. 1B) and ^77^STGGYG^82^ and ^114^GSSSQS^119^ from FUS. Four of these five structures were determined from X-ray data collected at the micro-focus beamline of the Advanced Photon Source (table S1), but to determine the structure of STGGYG we had to turn to micro electron diffraction (microED). For MicroED, we optimized the size of STGGYG crystals by gentle sonication and pipetting to achieve a crystal thickness of less than 1 μm (*19*) and data were collected to 1.1Å as previously described (*20*).

All five segments crystalized as pairs of β-sheets (Fig. 2). Each continuous β-sheet runs the length of the crystal, formed from the stacking of about 300,000 segments. Four of the structures show kinked pairs of sheets. The strands are kinked at either glycines or aromatic residues instead of being extended (fig. S2). Although these four structures differ in details, they share common adhesive features. Each segment hydrogen bonds in-register to an identical segment below it (Fig. 2A to D and fig. S3). Aromatic residues predominate in the kinked structures, both for inter-sheet stabilization and intra-sheet stabilization. Within sheets, the aromatic side-chains stack in an energetically favorable conformation, with the planes of the rings stacked parallel at a separation of 3.4Å(*21*–*23*) (fig. S3). These aromatic “ladders” enhance the stability of each β-sheet. In the four kinked structures, the kinks allow close approach of the backbones, providing favorable van der Waals or hydrogen bonding interactions between the sheets (fig. S3). These close interactions are reflected in the high values of the Structural Complementarity, Sc, shown in Fig. 2.

In contrast to the four kinked structures, that of GSSSQS is similar to the steric zippers formed by the adhesive segments of pathogenic amyloid proteins (*16*). That is, the β-strands are extended, mating at a tight, dry interface, with interdigitation of the sidechains extending from one sheet to its mate. Neighboring β-sheets mate in the kinked structures, but their kinks prevent side chains from interdigitating with each other across this β-sheet interface. Consequently the kinked interfaces bury smaller solvent-accessible surface areas than found in the interfaces in pathogenic amyloid fibrils, and presumably have lower binding energies. Because of the similarity of the kinked structures to pathogenic amyloid, we term these structures Low-complexity, Amyloid-like, Reversible, Kinked Segments or LARKS.

Calculations support our structural inference that LARKS have smaller binding energies than steric zippers. We estimated the energies of separation of the pairs of β-sheets in LARKS and also in steric zippers, by applying atomic solvation parameters (*24*, *25*) to our structures: the mean atomic solvation energy for separation of our LARKS interfaces is 393 ± 281 cal/mol/β-strand, whereas it is 1431 ± 685 cal/mol/β-strand for 75 steric zipper structures (fig. S4). While these estimates are crude, they suggest that the adhesive energy of one pair of β-strands in a LARKS is of the order of thermal energy at body temperature, so that pairs of β-sheets would adhere only through interactions of multiple pairs of strands. In contrast the adhesive energy of one pair of strands in a steric zipper is several times that of thermal energy. In short, the paired kinked β-sheets formed by segments of LC domain LARKS are less strongly bound to each other than are the β-sheets in pathogenic amyloid fibrils, yet still produce fibrils with the biophysical hallmarks of a cross β-diffraction pattern of pathogenic amyloid.

To identify potential LARKS in the human proteome, we used computational 3D profiling to thread sequences onto each of the atomic structures of SYSGYS, GYNGFG, and STGGYG (Fig. 3A) (*26*, *27*). In this procedure, we first removed side-chains from the structure of these three LARKS, producing three threading models. Then we threaded each human protein through these models, moving the sequence one residue at a time. For each segment of six residues on the threading model, we tested that segment for compatibility with its structure by computing the Rosetta energy (*28*) after optimally re-packing side-chains. That is, to thread an entire protein, each possible six-residue segment is threaded and scored in this manner to identify which segments of the protein can adopt a LARKS structure. In a preliminary test of this procedure, we threaded the sequences of the LC domains of FUS and hnRNPA1 on our structure of SYSGYS. From this we identified the segments GYNGFG and STGGYG as candidate LARKS; determining their structures confirmed them as LARKS (Fig. 2C and D). This validated our threading procedure, and encouraged us to apply the procedure to estimate the frequency of LARKS in the entire human proteome.

**Fig. 3.**
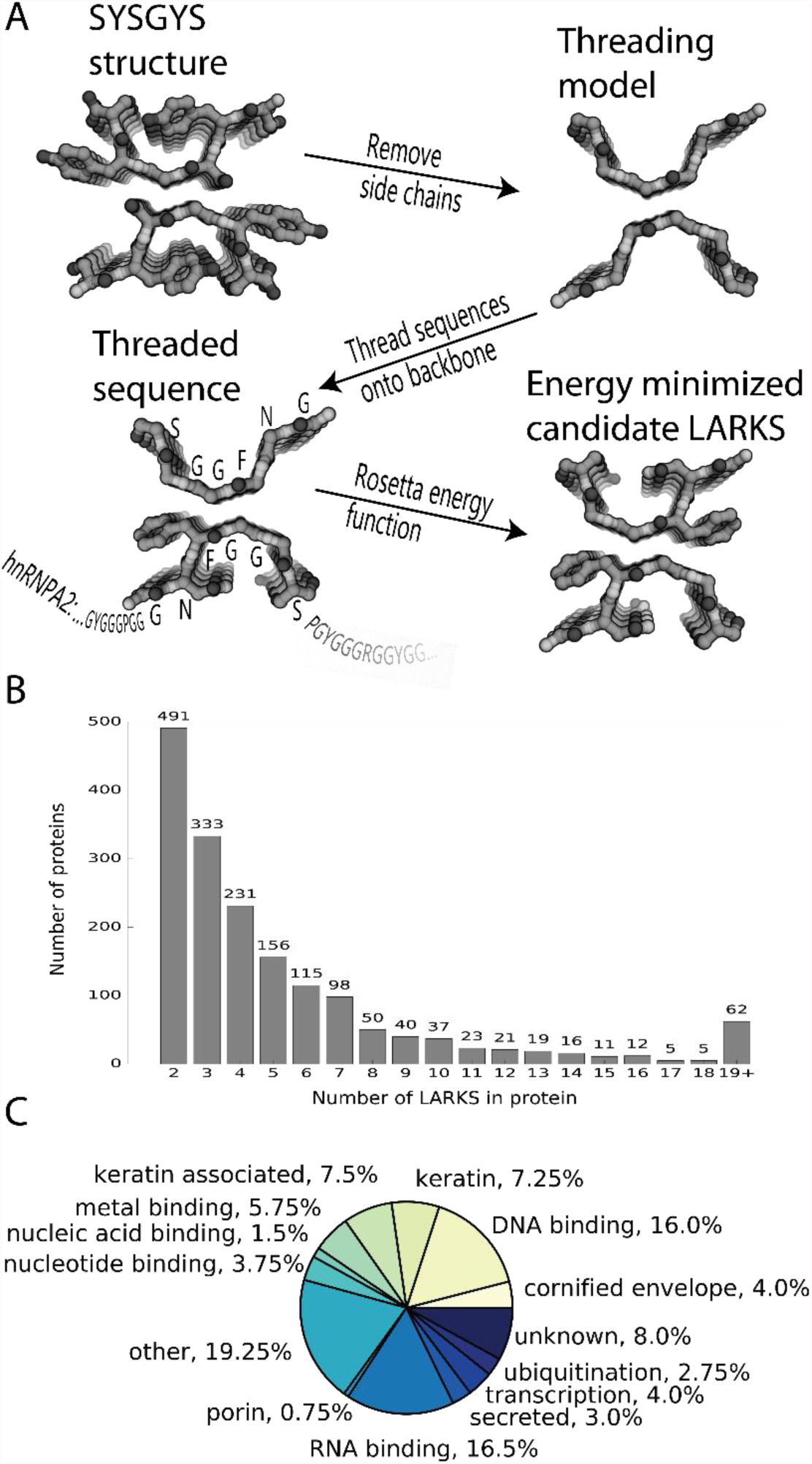
3D profiling to identify LARKS in LC domains of human proteins. **(A)** Method: Sidechains are removed from the backbones of one of our atomic structures of a LARKS. Then the sequence interest (hnRNPA1 shown) is threaded through the model, six residues at a time by placing the sidechains on the backbone of the model. Sidechains are repacked and a Rosetta energy function is used to estimate if the structure is favorable for the threaded sequence. **(B)** The frequency of the number of LARKS in 1725 human proteins predicted to house at least two LARKS. Proteins having two or more LARKS are predicted to have the capacity to form networks and possibly gels. **(C)** The annotated functions of the 400 proteins with the most predicted LARKS.

Analyzing the non-redundant human proteome of 20120 sequences from UniProt (see Materials and Methods), we found 5867 proteins with LC domains. Of these, 2500 proteins contain at least one LARKS and 1725 proteins contain two or more LARKS, thus having the capacity to form multivalent interactions. Hundreds of proteins house three or more LARKS (Fig. 3B). The 400 human LC domains most enriched in LARKS have an average of 14 LARKS each (fig. S5). Each of the three kinked backbones independently predicts most of the same proteins rich in LARKS (fig. S6). Further analysis (fig. S7) confirms that our profiling procedure identifies proteins enriched in a structural motif, rather than in a sequence motif.

We assigned cellular function to these 400 proteins based on their Uniprot annotations (Fig. 3C). Of the 400, 16% are DNA binding, 17% are RNA binding, and 4% are nucleotide binding, consistent with reports of nucleotide binding proteins being found in membraneless organelles (*7*, *8*). Unexpected to us, keratins (5%), keratin associated (9%), and cornified envelope proteins (4%) are also heavily enriched in LARKS. The finding of keratins is consistent with the experiments of Windoffer et al. (*29*) who showed that as keratin granules are trafficked to the cell cortex where they merge and eventually mature into filaments. The LC domains flanking the folded domains of the keratin may act as multivalent sites to allow keratin proteins to assemble into particles through LARKS and eventually allow the folded segments to form a keratin filament. Also rich in LARKS are proteins found in ribonucleoprotein particles such as the spliceosome or nucleolus (Fig. 4). Porin proteins such as nucleoporin nup54 with FG repeats are enriched in predicted LARKS, and previous literature shows that FG repeats purified in vitro form an amyloid-like hydrogel (*30*, *31*). This suggests that the FG repeats of nuclear porin may form LARKS as a basis for formation of the diffusion barrier of the pore.

**Fig. 4.**
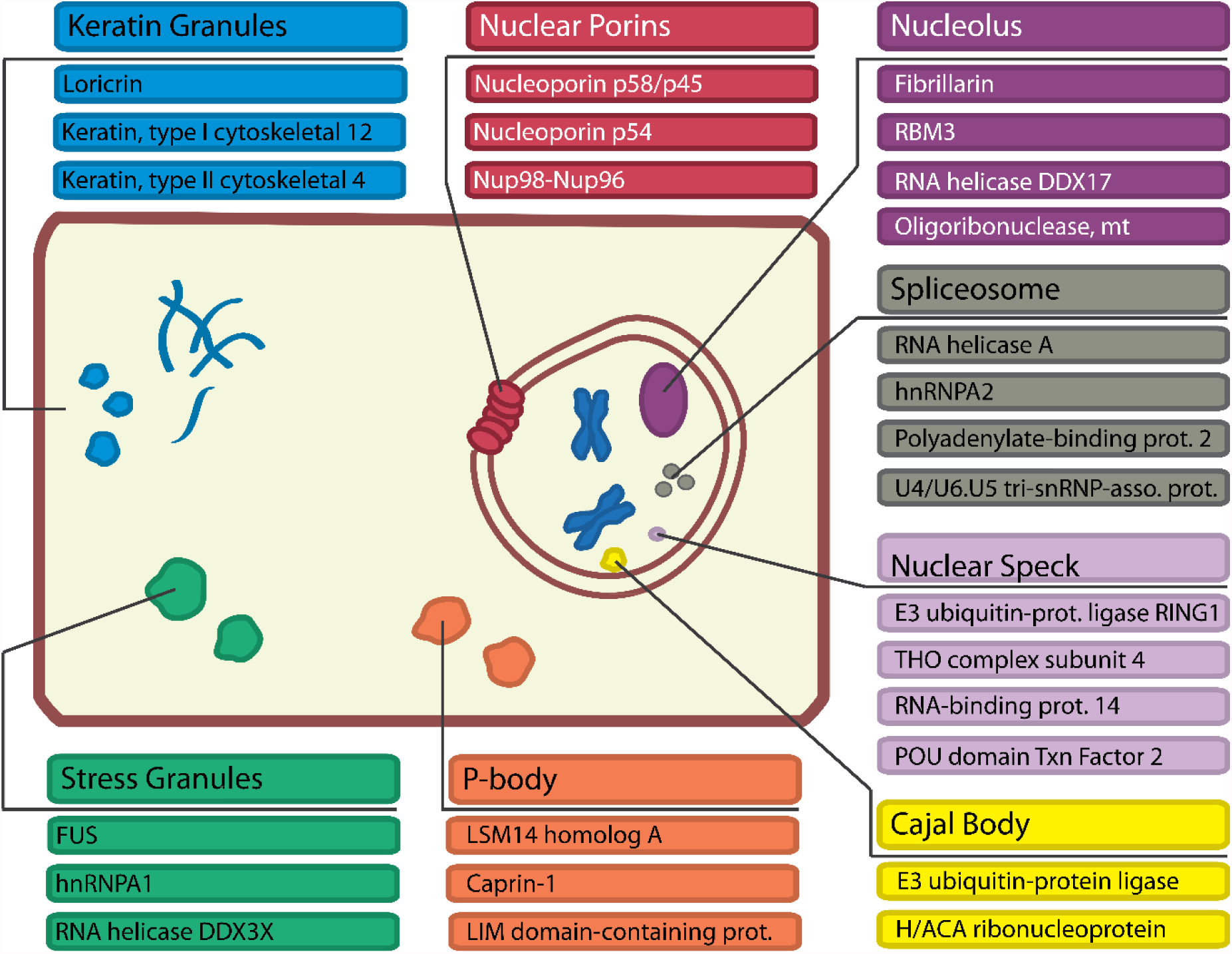
Functions of the 400 proteins most enriched in LARKS and dynamic intracellular bodies they are known to be a part of.

We assigned additional cellular functions to these 400 proteins from their associated gene ontology (GO) terms. We found certain GO terms enriched for these 400 proteins compared to the human proteome (Materials and Methods). GO terms related to RNA transport, processing, localization, and snoRNA are over represented (table S2). Proteins with the GO term for SG assembly are also enriched in LARKS. Epithelial cell differentiation was an enriched GO term due to the number of cornified envelope proteins and keratins enriched in LARKS. Therefore we propose 3D profiling for LARKS as a tool to identify proteins that may adhere by multivalent interactions, guiding compartmental organization in higher organisms in the form of membraneless organelles.

Our finding of numerous LARKS in the LC domains of the human proteome illuminates the function of these domains. Previous work has shown that RNA binding proteins and proteins found in SGs are enriched in LC domains, and that LC domains are sequestered into membraneless organelles such as SGs and nuclear paraspeckles (*3*, *4*, *8*). Therefore we propose that LARKS endow LC domains with multivalent interactions that can lead to phase separations and formation of dynamic intracellular bodies. We do not propose that the aromatic-dominated LARKS that we have observed are the only multivalent interactions that drive the formation of dynamic intracellular bodies. However, our GO analysis supports a connection between LARKS and these bodies.

The prevalence of LC domains within eukaryotic proteomes has long been recognized (*32*), but the role of these domains has not been fully defined. Previous discoveries include that: i) LC domains can “functionally aggregate” (*33*) ii) proteins with LC domains typically form more protein-protein interactions (*34*, *35*) iii) proteins can interact homotypically and heterotypically through LC domains (*3*, *4*, *36*).

Our atomic structures support the hypothesis that LC domains have the capacity to form gel-like networks and promote dynamic intracellular bodies. LARKS possess three properties that are consistent with their functioning as adhesive elements that enable the formation and dissolution of dynamic intracellular bodies: i) High aqueous solubility contributed by their high proportion of hydrophilic residues: serine, glutamine, asparagine, tyrosine; ii) High proportion of glycine ensures their flexibility; iii) Two or more interaction motifs per chain (fig. S5), endowing them with multivalency that enables them to entangle with multiple chains - in cis or trans - to facilitate entangling networks such as found in gels. That each LARKS provides adhesion only comparable to thermal energy suggests that numerous LARKS must cooperate in gel formation, and that the interactions must be highly concentration dependent and can be transient. If steric zippers act as molecular glue, then LARKS in LC domains act more akin to Velcro.

In conclusion, we propose LARKS as a protein interaction motif capable of providing sufficient adhesion to stabilize a variety of dynamic intracellular bodies. At the same time, LARKS seem sufficiently labile to permit reversible formation of these bodies as cellular conditions revert. LARKS add to the emerging paradigm that cells use LC domains to temporarily compartmentalize macromolecular complexes without the use of a membrane.

## Acknowledgments

Our x-ray diffraction data were collected at the Northeastern Collaborative Access Team beamline 24-ID-E which is funded by the National Institute of General Medical Sciences from the National Institutes of Health (P41 GM103403). This research used resources of the Advanced Photon Source, a U.S. Department of Energy (DOE) Office of Science User Facility operated for the DOE Office of Science by Argonne National Laboratory under Contract No. DE-AC02-06CH11357. We thank NSF MCB-1616265, NIH AG-054022, and HHMI for support.

## Supplementary Materials

### Materials and Methods

#### FUS purification and hydrogel formation

The plasmid with pHis-parallel-mCherry-1FUS214 was a generous gift from the McKnight Lab (UT Southwestern, Dallas, TX) and FUS was purified and formed into a gel according to the work presented in Kato et al. (*3*). A hydrogel was formed by placing purified mCherry-FUS concentrated to 100mg/ml in 4° for 2 months.

#### Negative stain EM

A pipette was used with remove 2μl of gel and then mix with 798μl H2O and pipetted vigorously to break up gel. This sample was pipetted on a copper mesh EM grid and stained with 2% uranyl acetate. The grid was scanned for fibrils and images were collected on a T12 120kV electron microscope (FEI) with a Gatan 2kX2k CCD.

#### Protein segment crystallization and structure determination

FUS | 37-SYSGYS-42 | peptide was ordered from GenScript (Piscataway, NJ). A peptide stock solution was prepared by adding nano-pure H2O to achieve a concentration of 10mg/ml. Solubility was enhanced by autoclaving, and then the peptide solution was immediately used for crystallization. Crystals grew at room temperature by hanging drop vapor diffusion. The reservoir contained 0.1 M Bis-Tris pH 6.0, 0.2 M magnesium formate dihydrate. Crystals were cryoprotected by transferring them into 50% glycerol and then mounted on a loop. Diffracton data were collected at the Advanced Photon Source (APS) on beamline 24-ID-E at 100 K. Phases were determined by molecular replacement using the program phaser (*37*). The successful search model was a truncated form of the peptide MIHFGND (PDB ID code 3NVH) with sidechains and methionine removed. Refinement was performed with Refmac (*38*).

FUS | 54-SYSSYGQS-61 | peptide was ordered from GenScript (Piscataway, NJ). The peptide was solubilized by adding nano-pure H2O to achieve a concentration of 75mg/ml, and then autoclaved. The peptide solution was immediately used for crystallization. Crystals grew at room temperature by hanging drop vapor diffusion. The reservoir contained 0.5 M ammonium tartrate dibasic pH 7.0. Crystals were cryoprotected by transferring them into 50% glycerol and then mounted on a loop. Diffraction data were collected at the APS beamline 24-ID-E at 100 K. Phases were determined by direct methods using the program ShelxD (*39*). Refinement was performed with Refmac (*38*).

FUS | 114-GSSSQS-119 | peptide was ordered from Innopep (San Dieo, CA). The peptide was dissolved in nano-pure H2O at 150mg/ml. Crystals grew at room temperature by hanging drop vapor diffusion. The reservoir contained 0.1 M citric acid pH 4.0 and 2.4 M ammonium sulfate. Crystals were cryoprotected by transferring them into 50% glycerol and then mounted on a loop. Diffraction data were collected at the APS beamline 24-ID-E at 100 K. Phases were determined by molecular replacement using the program phaser (*37*). The successful search model was an idealized poly alanine β-strand. Refinement was performed with Refmac (*38*).

hnRNPA1 | 243-GYNGFG-248 | peptide was ordered from Innopep (San Dieo, CA). The peptide was solubilized by adding nano-pure H2O to achieve a concentration of 10mg/ml, and then autoclaved. The peptide solution was immediately used for crystallization. The crystals grew at room temperature by hanging drop vapor diffusion. The reservoir contained 0.2 M magnesium acetate tetrahydrate, 0.1 M sodium cacodylate pH 6.5, and 30% (v/v) MPD. Crystals were mounted dry on the ends of pulled glass needles. Data were collected at the APS beamline 24- ID-E at 100 K. Phases were determined by direct methods using the program ShelxD (*39*). Refinement was performed with Refmac (*38*).

All X-ray diffraction data sets were collected on an ADSC Q315 CCD detector. All diffraction data were processed with XDS (*40*).

FUS | 77-STGGYG-82 | peptide was ordered from Innopep (San Dieo, CA). The peptide was dissolved in nano-pure H_2_O at 150mg/ml. Crystals grew at room temperature by hanging drop vapor diffusion. The reservoir contained 0.1 M sodium acetate pH 4.6, 0.15 M ammonium sulfate, and 25% (v/v) PEG 2000 MME. Several drops were then collected using a pipette and put in a 200μl PCR tube and placed in a water bath sonicator. The solution was sonicated 20min with ice to prevent melting the peptide crystals by temperature increase. A 2-3μl drop of resulting suspension was deposited onto a Quantifoil holey-carbon EM grid. Grids were then blotted and vitrified by plunging into liquid ethane using a Vitrobot Mark IV (FEI). On a per-crystal basis, blotting times and forces were optimized to keep a desired concentration of crystals on the grid and to avoid damaging the crystals. Frozen grids were then either immediately transferred to liquid nitrogen for storage or placed into a Gatan 626 cryo-holder for imaging. Images and diffraction patterns were collected from crystals using an FEG-equipped FEI Tecnai F20 TEM operating at 200 keV. All data were recorded using a bottom mount TVIPS F416 CMOS camera with a sensor size of 4000 by 4000 pixels, each 15.6 by 15.6 μm in size. Diffraction patterns were recorded by operating the detector in a continuous capture mode termed ‘rolling shutter’ with 2 × 2 pixel binning. This produced images 2000 by 2000 pixels in size. Exposure times for these images ranged from 1-2 seconds per image. During each exposure, crystals were unidirectionally rotated within the electron beam at a fixed number of degrees per second, corresponding to a fixed angular wedge per frame.

Crystals that appeared undistorted and that were 100-300nm thick produced the best diffraction. Since our crystals are needle-like and lie preferentially oriented on EM grids, data sets were collected from multiple crystals in different orientations and these sets were merged in order to achieve sufficient data completeness and redundancy. Data from individual crystals spanned a wedge of reciprocal space ranging from 60-80°, within 100-200 frames. We used a selected area aperture with an illuminating spot size of approximately one micron. The geometry detailed above equates to an electron dose of less than 0.1 e−/Å^2^ per second being deposited onto our crystals. Measured diffraction images were converted from tiff or tvips format into SMV crystallographic format, using in-house software available for download at https://cryoem.janelia.org/downloads/ (*41*).

Electron diffraction images were indexed and integrated with XDS (*40*) and data sets originating from different crystals were scaled together with XSCALE (*40*). Phases for FUS-STGGYG were determined by direct methods using the program ShelxD (*39*). Crystallographic refinement were performed using phenix (*42*) and buster (*43*) and a library of electron scattering factors.

All models were built with Coot (*44*).

#### Crystal powder diffraction

All images were collected from crystal clusters or hydrogel formed according to above protocols. All samples were shot wet and without cryo-stream. Images were collected with a RIGAKU R-AXIS HTC imaging plate detector using Cu K(alpha) radiation from a FRE+ rotating anode generator with VARIMAX HR confocal optics (Rigaku, Tokyo, Japan). The sample-to-detector distance was 200mm. The sample rotated 5° during the exposures.

#### Atomic solvation energy calculations

For each structure, crystal symmetry was used to generate a fibril model with two β-sheets each composed of 10 strands. The exposed surface area for each atom was measured with a 1.4Å diameter probe using the program areaimol from the CCP4 package (*45*). One β-sheet was translated away and the exposed surface area was measured again. These values were subtracted to find the difference in exposed surface area for each atom (Δ*A_i_*). The exposed surface area was then multiplied by an atomic solvation parameter (*24*) to get the estimated free energy for solvating that atom (*σ_i_*).

Atomic solvation parameters (*σ_i_*) for each atom:

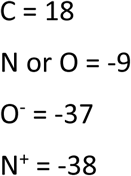

All atom atomic solvation energies were then summed for a strand:

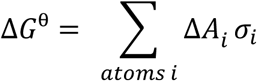

N-terminal and C-terminal charges on the peptide backbone were considered neutral, as they would be in the full length protein. The atomic solvation energy then reflects the calculated energy to melt the fibril interface. Energies are taken from a single, most central strand in one β-sheet.

#### Identifying LC domains

To identify LC domains on proteins we used the SEG algorithm (*2*) with normal parameters. We looked for stretches of at least 25 adjacent residues to consider the region to be an LC domain, but allowed for an interruption by up to five residues not considered LC to get rid of artifacts in the algorithm. This allowed us to identify 178 proteins with LC domains out of a UniProt proteome of 3990 proteins in *Escherichia coli*; 1202 proteins with LC domains out of a UniProt proteome 6720 proteins in *Saccharomyces cerevisae*; and 5893 proteins with LC domains out of a UniProt proteome of 20197 proteins in *Homo sapiens*. Minimal proteomes were downloaded from the UniProt consortium (*46*): *Homo sapiens* was downloaded March 28, 2016; *Escherichia coli* was downloaded April 2, 2016, and *Saccharomyces cerevisae* was downloaded April 4, 2016.

#### Template based threading

Using a process analogous to our 3D profile method for predicting steric-zipper-forming segments (*27*), we used RosettaDesign for a high-throughput computational search of other sequences compatible with the STGGYS, SYSGYS, and GYNGFG backbones. Each structural backbone template consists of the assembly of two (STGGYS, GYNGFG) or three (SYSGYS) beta-sheets, with each sheet made up of ten hexapeptides with the same residue sequence. Method for identifying interface for STGGYG structure is shown in Fig. S8. We threaded the sequences of 20,120 proteins from a non-redundant human proteome (*46*) that collectively contains 7,900,599 unique 6-residue segments. Segments were evaluated with the standard Rosetta energy function (Talaris2014) except with changes to the Dunbrack rotamer and Lennard-Jones potential terms. For the Dunbrack term, we increased by a factor of 4 the acceptable standard deviation in the chi-angle. To de-emphasize repulsion, we increased the lj_switch_dis2sigma distance from 0.6 to 0.8 Angstroms. Models with energies below: -3.0 for NKGAII, 4.0 for STGGYG, 10.0 for GYNGFG, and -3.0 for SYSGYS (modified Rosetta Energy Units) were considered compatible with these structure backbones. Energy cutoffs for the steric zipper NKGAII were determined as in Goldschmidt et al. (*27*). Energy cutoffs for LARKS were determined by looking at a histogram of scores and choosing the most energetically favorable segments until the second inflection point of the curve. We determined the fraction of compatible segments in each protein. If any segment scored well on any of the three LARKS backbones, and it was in a LC domain, it was considered favorable for forming a LARKS. We only considered LARKS in LC domains because in globular domains, it would be unlikely that they would be available for interaction.

#### Identifying top 400 proteins enriched in LARKS

LARKS were found using the above methods. If at least six adjacent residues were predicted to be in LARKS, it was scored as a single LARKS. Proteins were ordered by the largest number of LARKS, and the top 400 were qualitatively categorized (Fig. 3C) based upon their UniProt GO annotations. For more rigorous methodology we used GO to find enriched GO terms in these top 400 proteins.

#### Gene ontology on top 400 proteins enriched in LARKS

The gene ontology tree and annotations for the human proteome were downloaded from the Gene Ontology Consortium (*47*). The GO terms for the top 400 proteins were found 4412 unique GO terms represented by these top 400 proteins. We used standard bootstrapping methods to identify statistically enriched terms. For bootstrapping, to test if a GO term was enriched, 400 random proteins were chosen from the human proteome and we counted how many times that GO term was found in those 400 proteins. This procedure was repeated 15,000 times to create a distribution of the number of times that GO term was found in a random selection from the proteome. If the actual number of times that GO term was found in the top 400 proteins rich in LARKS was greater than the 95^th^ percentile of times that GO term was found in our random distribution, then it was considered enriched. P-values are from the score on the randomly selected distribution. This yielded 496 enriched GO terms. Several were qualitatively selected for display in supplemental table S2.

#### Venn diagram overlap of proteins rich in structures predicted by different threading backbones

For fig. S6 we threaded the human proteome on four different backbones: SYSGYS, STGGYG, GYNGFG, and NKGAII. The first three being examples of kinked backbones to search for LARKS and the latter being a steric zipper to search for traditional amyloid forming segments. For each peptide backbone, all proteins in the human proteome were ranked by number of hits for that backbone, and then the top 400 were taken. Overlap of proteins predicted by each backbone was found, and presented on a 4-way Venn diagram. Low-complexity domains were not considered in determining if a segment was a hit, as in prediction for LARKS in the rest of the paper.

#### Venn diagram of sequence bias and predicted structures

For fig. S7, we ranked the human proteome according to these four methods:

1. Highest fraction on the protein in predicted LC domains (as described in Materials and Methods)
2. Highest content of Glycine or Serine (these residues are highly over represented in human LC domains)
3. Most number of predicted LARKS (as described in Materials and Methods)
4. Most number of predicted steric zippers (disregards LC domains)

Then the top 400 proteins for each category were taken and overlap between the 4 methods was represented on a 4-way Venn diagram.

#### Steric zipper model from GSSSQS

To make an extended model of the steric zipper GSSSQS, we made a pseudo symmetry related GSSSQS model and in Coot, moved the new 6-residue segment to the N or C term of the original models. The chains were joined and idealized Phi-Psi angles were used from Coot to idealize the beta-strands. Mutations were introduced in Coot and by choosing the appropriate rotamer. The mutation showing a kink was made by joining a model of GSSSQS with a model of GYNGFG using the same methodology as above. Images were made using PyMol.

## Supplementary Figures

**Fig. S1.**
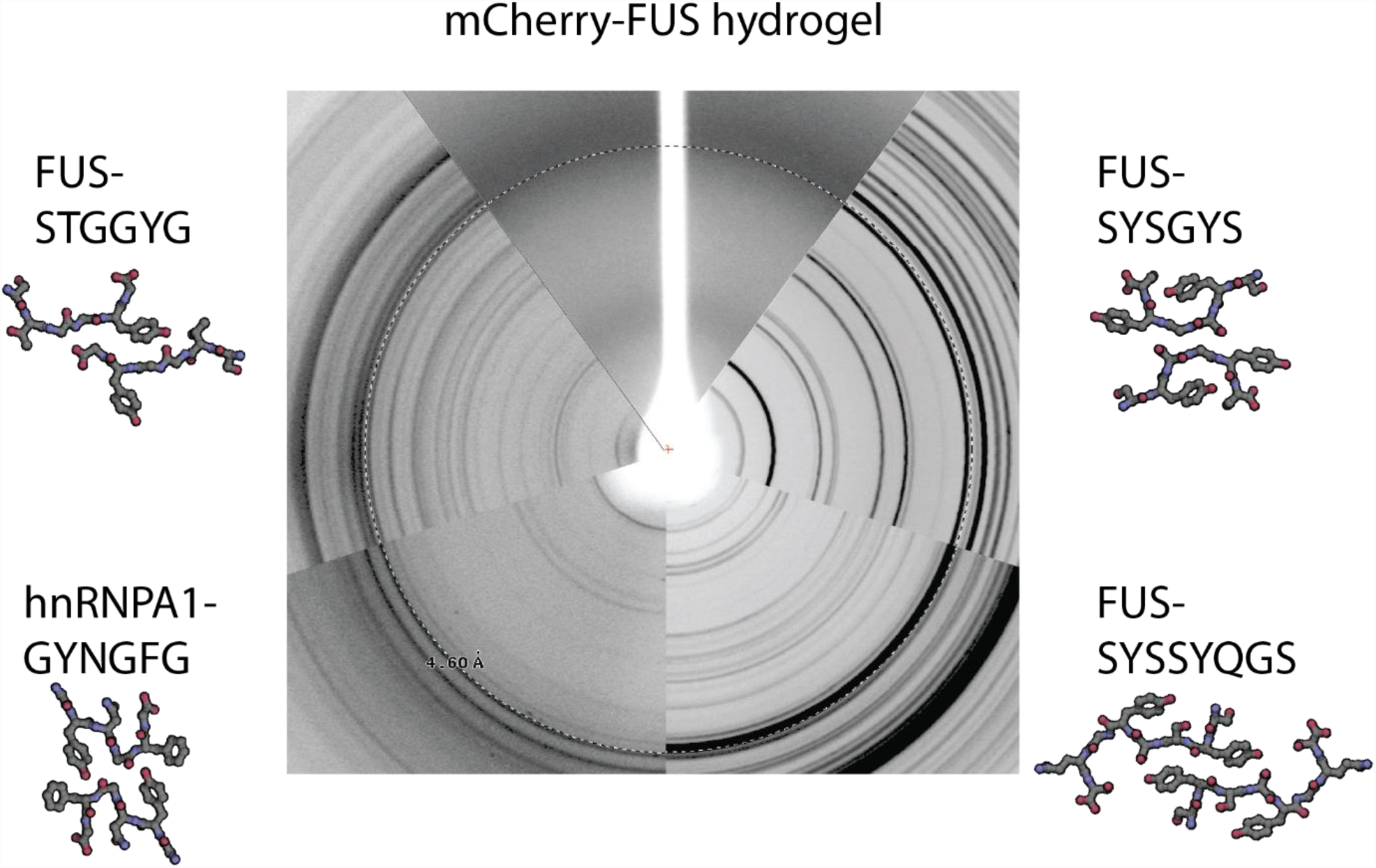
Fiber X-ray diffraction of LARKS compared to that of FUS hydrogel. Each wedge shows the fiber diffraction of the corresponding samples recorded under identical conditions (distance to detector, X-ray wavelength, and detector) to allow direct comparison of Bragg spacings. The patterns from the four LARKS are the crystalline powder diffraction patterns. The dotted circle shows spacing of 4.60 Å. All LARKS display a ring at, or very near, 4.60 Å indicating their consistency with the diffraction pattern from the FUS LC domain.

**Fig. S2.**
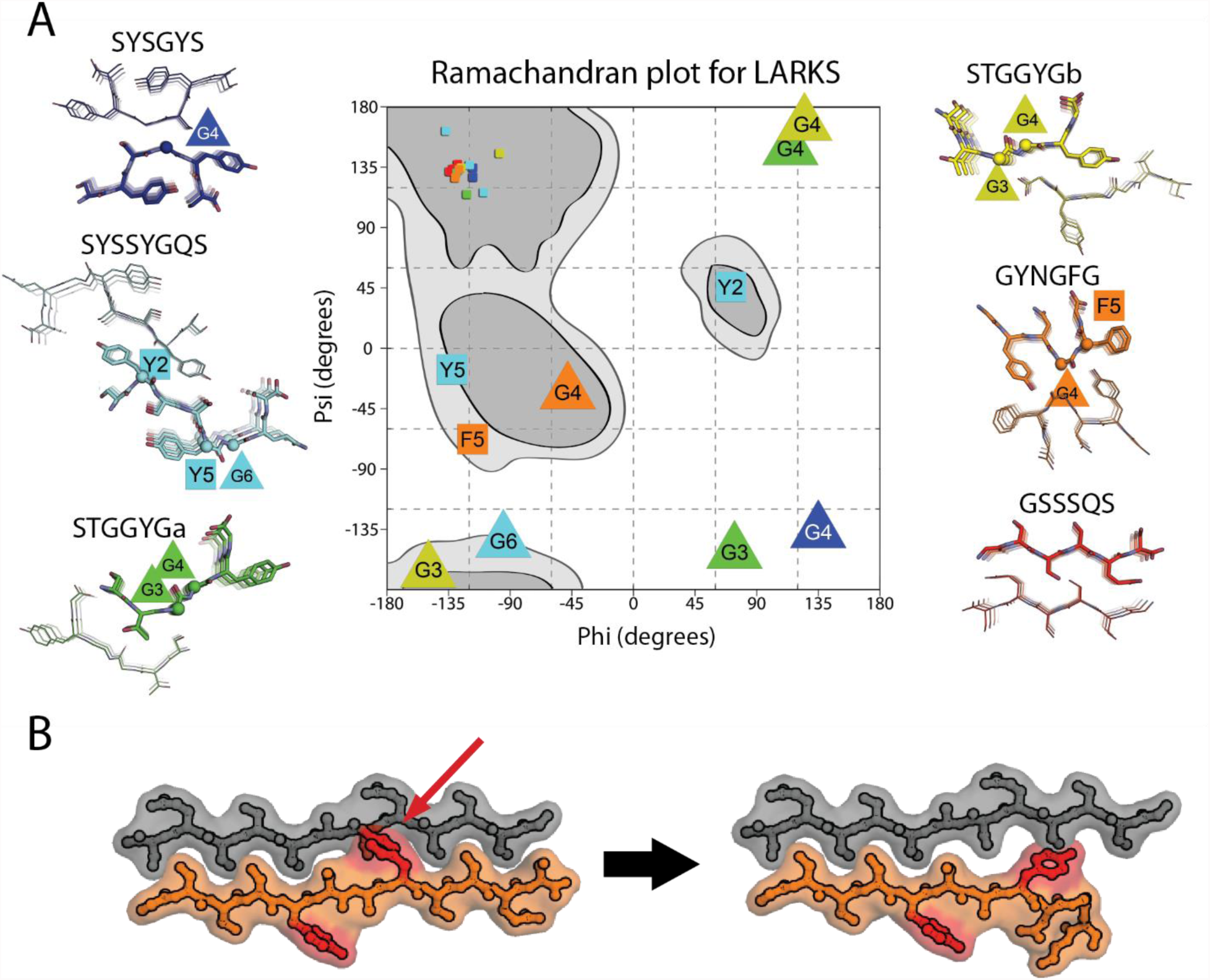
The structural origin of kinks in LARKS. **(A)** Dihedral (Phi-Psi) angles for all residues in our atomic structures of LARKS and GSSSQS, a steric zipper, shown in a Ramachandran diagram. The Phi-Psi angles are color coded by structure. Large triangles indicate glycine residues that occupy non-beta Phi-Psi angles, and large squares represent aromatic residues that occupy non-beta Phi-Psi angles. Phi-Psi angles fall in allowed regions of the Ramachandran diagram. **(B)** Models of a steric zipper based on our structure of GSSSQS with tyrosine residues replaced at positions I and I+3; these produce a steric clash with the β-strand in the opposing sheet (red arrow). To accommodate the tyrosine in the mating interface, the peptide backbone of the orange strand must bend away from the gray strand, producing the kink.

**Supplemental Figure 3.**
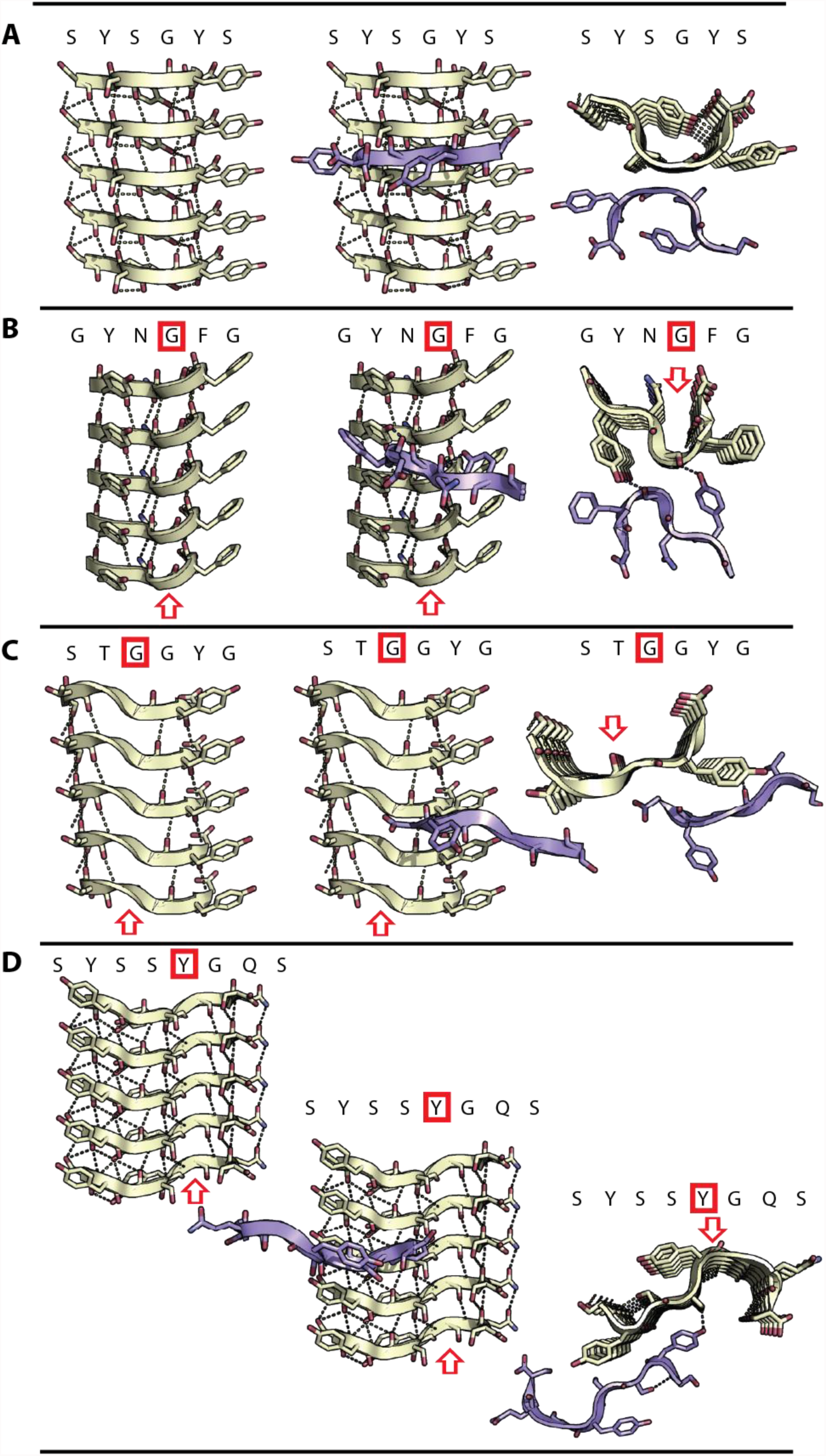
Intra-sheet and inter-sheet patters of hydrogen bonding in four LARKS. The left-hand column views five stacked beta-strands in our four LARKS viewed perpendicular to their fibril axes with the corresponding segment sequence above. The second column displays one strand of the mating sheet, colored blue, in the foreground. The third column views the fibril parallel to the fibril axis. Intra-sheet hydrogen bonds are shown by yellow dots; inter-sheet hydrogen bonding from unsatisfied backbone donors/acceptors are in purple. Red boxes on the sequence indicate residues with carbonyl/amide backbone hydrogen bond donors/acceptors that are not satisfied by hydrogen bonding along the fibril axis, and instead form hydrogen bonds with mating sheets. Red arrows point to hydrogen bonds unsatisfied by secondary structure. **(A)** Structure of SYSGYS **(B)** structure of GYNGFG **(C)** structure of STGGYG **(D)** structure of SYSSYGQS.

**Fig. S4.**
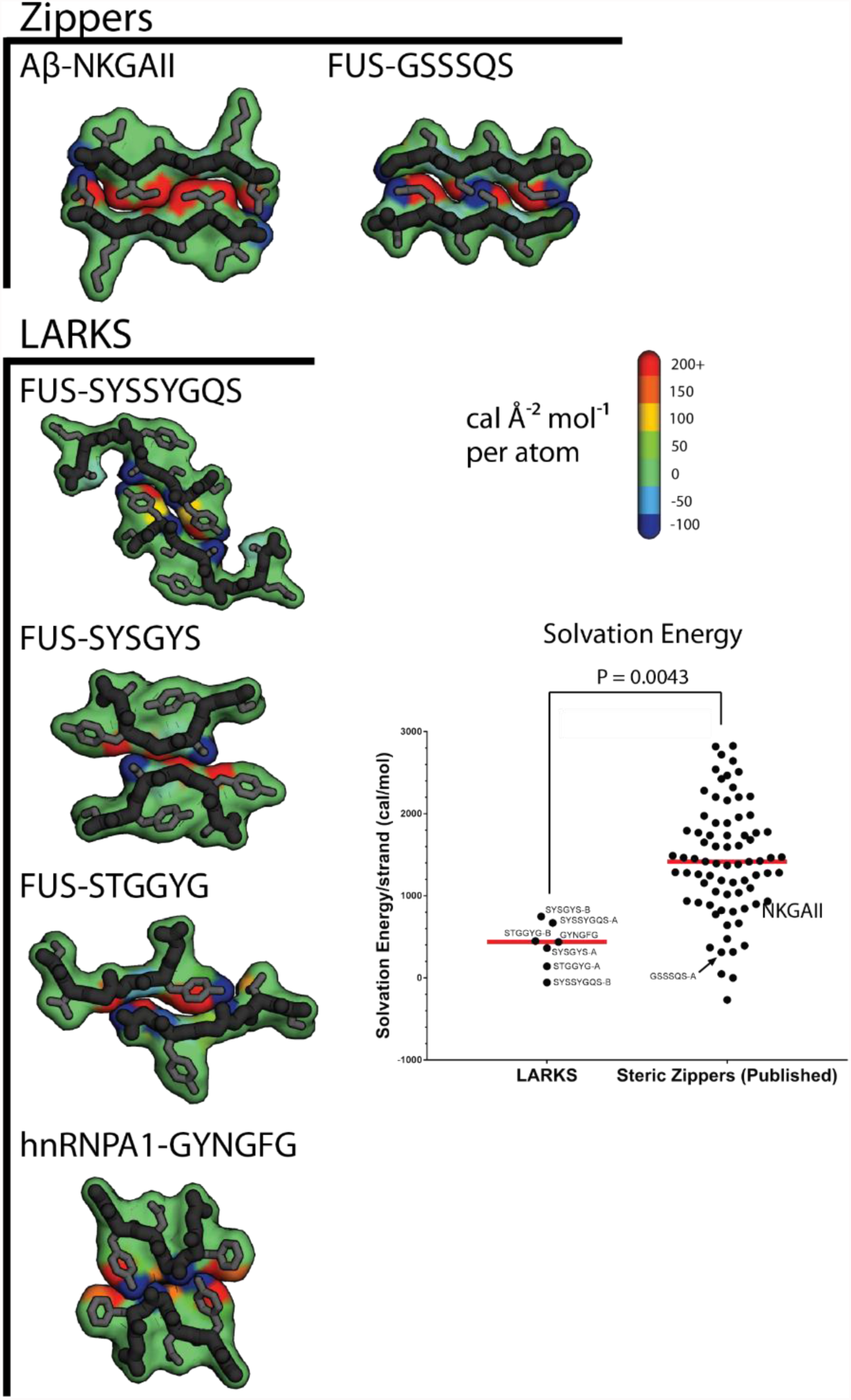
Solvation energies for LARKS and steric zippers. We calculated solvation energies for the process of separating (expose to solvent) two mated sheets. Values are in calories per mol per for the buried surface area. Plot compares the solvation energy for four LARKS published in this work and 76 published steric zipper structures. Steric zippers have significantly more stable interfaces as judged by these energy values (P = 0.0043 by student’s t-test). GSSSQS has an unusually low solvation energy for a steric zipper because most of the atoms lining the interface are hydrophilic (side chain hydroxyl oxygen atoms).

**Fig. S5.**
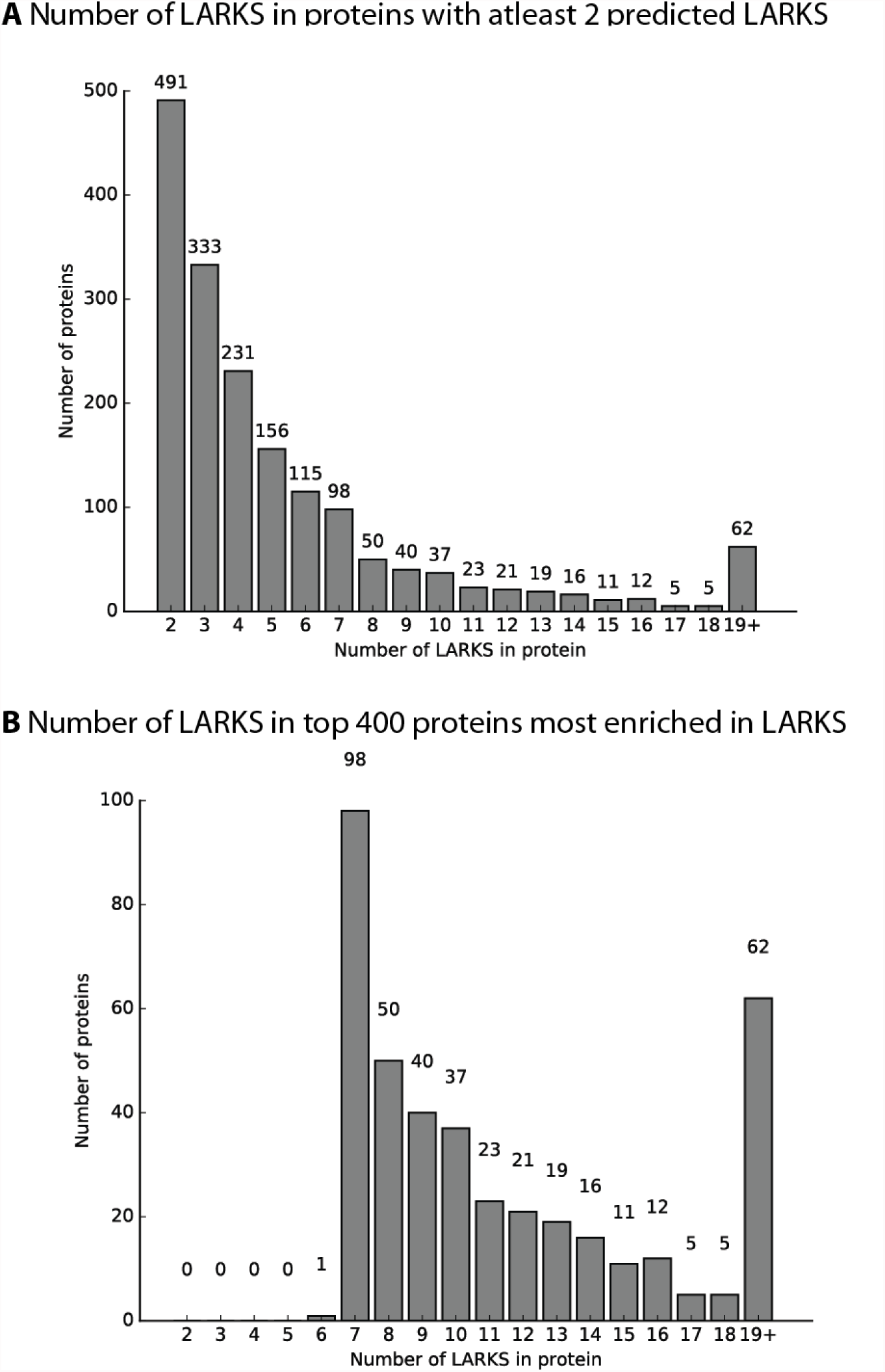
Histogram with the frequency of predicted LARKS in proteins in the human proteome. **(A)** The frequency of the number of LARKS in 1725 human proteins (of 20120 threaded proteins) predicted to house at least two LARKS. Proteins having two or more LARKS are predicted to have the capacity to form networks and possibly gels. **(B)** The frequency of LARKS in the top 400 proteins most enriched in LARKS.

**Fig. S6.**
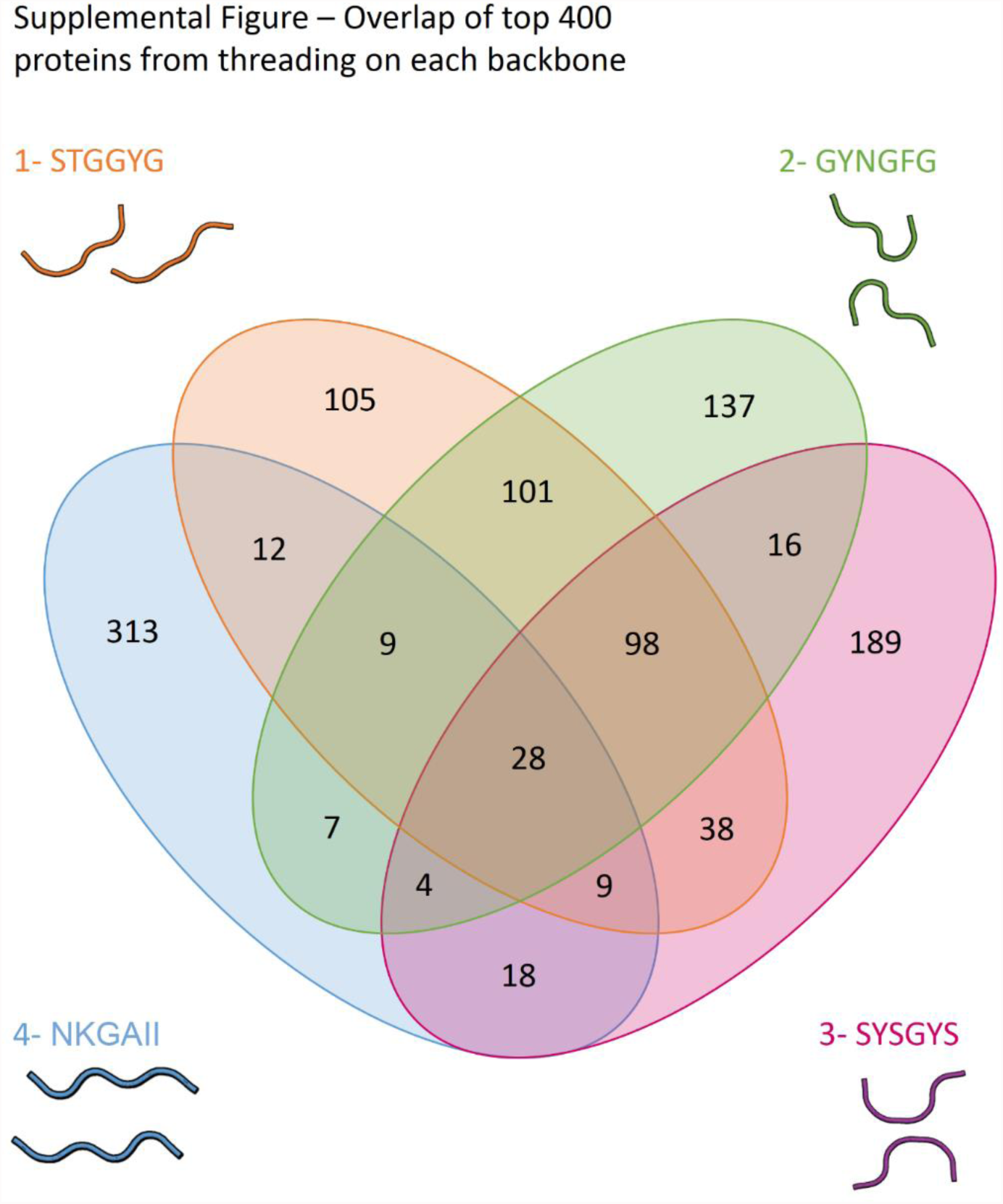
3D profiling on three different LARKS backbones largely identify the same proteins as rich in LARKS. The top 400 proteins rich in LARKS as judged by treading the human proteome on backbones: 1) STGGYG, 2) GYNGFG, 3) SYSGYS, or 4) NKGAII – a steric zipper for reference. 124 proteins are agreed to be the top 400 proteins enriched in LARKS by each different LARKS backbone, indicating that each method of threading largely agrees with the others. 384 proteins are unique to the steric zipper backbone, indicating that the steric zippers are distinct structural motifs from LARKS.

**Fig. S7.**
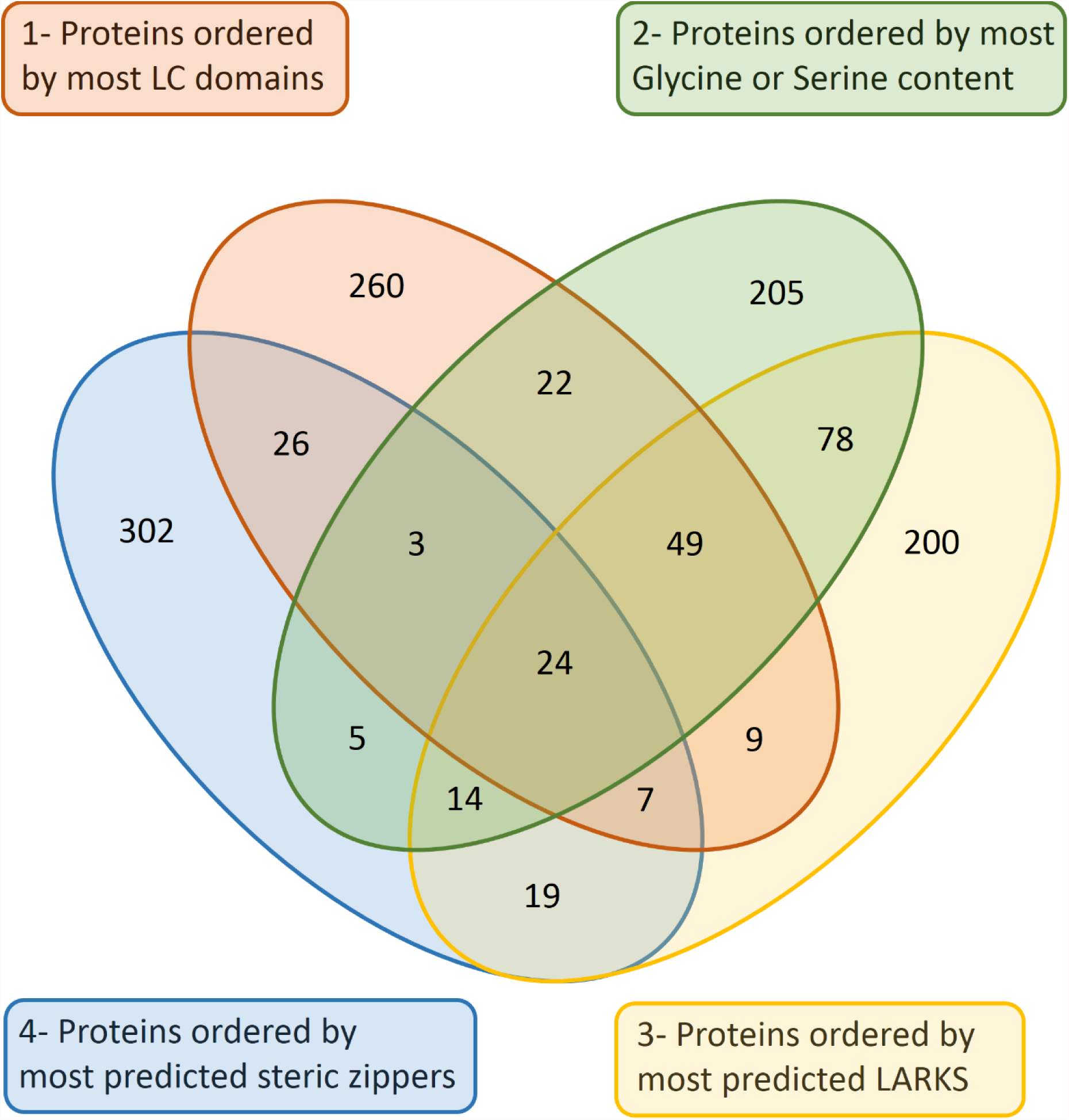
3D profiling identifies a unique set of LARKS-containing proteins, which are not simply LC-containing proteins or serine and glycine containing proteins. We compared the identities of the top 400 proteins enriched in predicted LARKS (yellow oval) to the top 400 proteins by fraction of residues in steric zippers (blue oval), in LC domains (orange oval), or by serine/glycine sequence bias - some of the most over-represented amino acids in LC domains in humans (green oval). The lack of large overlap between the circles indicates that that 3D profiling identifies a relatively unique set of 400 proteins, which are not selected by LC domains, by sequences rich in serine/glycine, or rich in steric zippers. In other words, 3D profiling selects proteins by 3D structure and not by sequence.

**Fig. S8.**
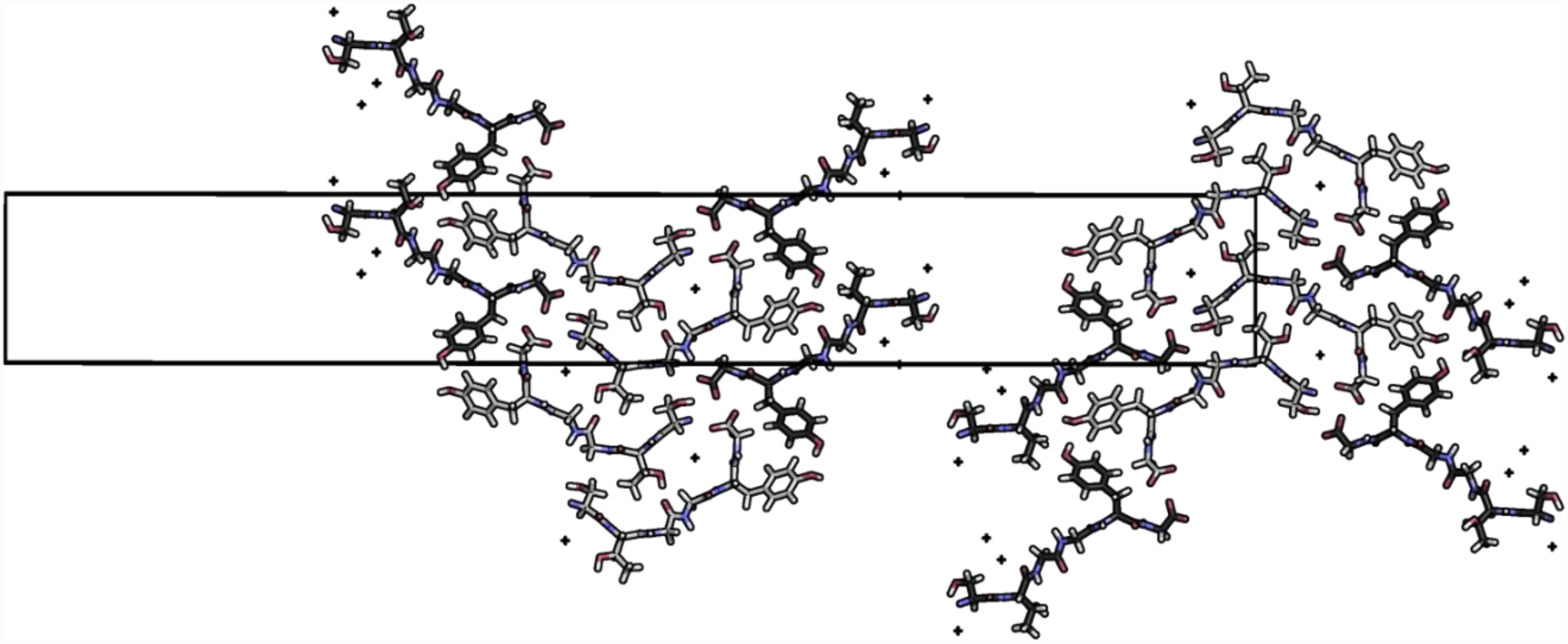
Crystal structure of STGGYG determined by microED reveals an asymmetric unit with two segments of STGGYG in different conformations. The unit cell outlined by the black box. Nitrogen atoms are blue; oxygens are red red; waters are represented by small crosses; carbons of STGGYGa are light gray; and carbons of STGGYGb are colored dark gray. Two paired interfaces between STGGYG segments exist in the crystal structure: an interface between STGGYGa and STGGYGa, and an interface between STGGYGa and STGGYGb. We used the interface between STGGYGa and STGGYGb as a LARKS in our 3D profiling because it has the more extensive and hydrophobic interface.

**Table S1.** Crystallographic statistics. Statistics about data collection and structure determination for new structures presented in this work. MR = Molecular Replacement; DM = Direct Methods; * = includes hydrogen atoms.

| Peptide | SYSGYS | SYSSYGQS | STGGYG | GSSSQS | GYNGFG |
| --- | --- | --- | --- | --- | --- |
| Protein | FUS | FUS | FUS | FUS | hnRNPA1 |
| Crystal size (in $\mu\text{m}$) | 150 x 6 x 6 | 200 x 8 x 8 | 1 x 0.2 x 0.2 | 150 x 7 x 7 | 200 x 7 x 7 |
| Data Collection |  |  |  |  |  |
| Diffraction type | X-ray | X-ray | Electron | X-ray | X-ray |
| Beamline | APS 24-ID-E | APS 24-ID-E | JFRC TF20 | APS 24-ID-E | APS 24-ID-E |
| Space group | P 1 21 1 | C 1 2 1 | P 21 21 21 | C 1 2 1 | P 21 21 21 |
| Resolution ($\text{\AA}$) | 1.1 | 1.1 | 1.1 | 1.1 | 1.1 |
| Unit cell dimensions: a, b, c ($\text{\AA}$) | 17.57, 4.82, 18.28 | 49.58, 4.79, 21.81 | 13.79, 4.93, 101.9 | 43.94, 4.80, 11.87 | 16.61, 4.77, 40.97 |
| Unit cell angles: $\alpha, \beta, \gamma$($^\circ$) | 90.00, 91.03, 90.00 | 90.00, 95.46, 90.00 | 90.00, 90.00, 90.00 | 90.00, 93.62, 90.00 | 90.00, 90.00, 90.00 |
| Molecules per asymmetric unit | 1 | 1 | 2 | 1 | 1 |
| Measured reflections | 5518 | 18360 | 23,271 | 3243 | 32383 |
| Unique reflections | 1176 | 2166 | 3220 | 1084 | 1584 |
| Completeness (%) | 82.0 (28.4) | 92.8 (83.3) | 95.5(74.9) | 95.0 (91.3) | 81.0 (36.5) |
| Redundancy | 4.7 (2.2) | 8.5 (4.4) | 7.2(3.9) | 3 (2.2) | 25.9 (6.8) |
| R<sub>merge</sub> | 0.179 (0.468) | 0.135 (0.547) | 0.250(0.553) | 0.217 (0.501) | 0.130 (0.250) |
| I/$\sigma$ | 7.8 (2.4) | 15.0 (4.0) | 4.15(1.5) | 5.39 (1.61) | 21.1 (8.7) |
| CC<sub>1/2</sub> | 0.989 (0.497) | 0.995 (0.800) | 0.985 (0.729) | 0.987(0.836) | 0.998 (0.962) |
| Refinement |  |  |  |  |  |
| Phasing method | MR | DM | DM | MR | DM |
| Rwork | 0.148 | 13.7 | 0.219 | 0.156 | 0.061 |
| Rfree | 0.176 | 14.8 | 0.255 | 0.176 | 0.083 |
| RMSD bond length ($\text{\AA}$) | 0.03 | 0.004 | 0.01 | 0.01 | 0.018 |
| RMSD angle ($^\circ$) | 2.27 | 0.755 | 1.06 | 1.39 | 2.22 |
| Number of peptide atoms* | 94 | 124 | 134 | 76 | 44 |
| Number of solvent atoms | 2 | 24 | 14 | 2 | 4 |
| Average B factor of peptide ($\text{\AA}^2$) | 4.2 | 8.3 | 7.4 | 3.9 | 1.4 |
| Average B factor of solvent ($\text{\AA}^2$) | 6 | 20.1 | 14.6 | 6.6 | 11.7 |
| PDB ID code |  |  |  |  |  |

**Table S2.** Selected enriched GO terms in the top 400 proteins rich in LARKS. The top 400 proteins rich in LARKS were analyzed for enriched GO terms. 496 GO terms were found to have P values below log10 -1.301 (equivalent to a p-value of 0.05). The first column is GO ID, the second column gives the name of the GO term. The P-value is the confidence of enrichment determined by standard bootstrapping methods.

| GO term ID | GO term description | log10 p-value |
| --- | --- | --- |
| GO:0051028 | mRNA transport | -3.875 |
| GO:0006397 | mRNA processing | -4.176 |
| GO:0030855 | epithelial cell differentiation | -4.176 |
| GO:0071765 | nuclear inner membrane organization | -1.689 |
| GO:0034063 | stress granule assembly | -1.860 |
| GO:0006403 | RNA localization | -3.398 |
| GO:0007004 | telomere maintenance via telomerase | -2.257 |
| GO:0016074 | snoRNA metabolic process | -2.745 |

## Any Additional Author notes

### Author contributions

DSE and MPH conceived of the project; MPH, MRS, JAR, and DC collected diffraction data and determined structures; MPH, MRS, and LG carried out computations; and MPH, MRS, and DSE wrote the manuscript with contributions from all authors.

